# The Strongest Protein Binder is Surprisingly Labile

**DOI:** 10.1101/2024.02.18.580677

**Authors:** Alba Fernandez-Calvo, Antonio Reifs, Laura Saa, Aitziber L. Cortajarena, David De Sancho, Raul Perez-Jimenez

## Abstract

Bacterial adhesins are cell-surface proteins that anchor to the cell wall of the host, thus initiating infection. The initial step in infection is precisely the binding to fibrinogen (Fg) from human tissue, after which bacteria can colonize the heart valves by the formation of biofilms. The study of this family of proteins is hence essential to develop new strategies to fight bacterial infections. In the case of *Staphylococcus aureus*, there exists a type of adhesins known as Microbial Surface Components Recognizing Adhesive Matrix Molecules (MSCRAMMs). Here, we focus on one of them, the Clumping Factor A (ClfA), which has been found to bind Fg through the dock-lock-latch (DLL) mechanism. Interestingly, it has recently been discovered that MSCRAMMs proteins employ a catch-bond to withstand forces exceeding 2 nN, making this type of interaction as mechanically strong as a covalent bond. However, whether this strength is an evolved feature characteristic of the bacterial protein or is typical only of the interaction with its partner is not known. Here we combine single-molecule force spectroscopy (smFS), biophysical binding assays and molecular simulations to study the intrinsic mechanical strength of ClfA. We find that despite the extremely high forces required to break its interactions with Fg, ClfA is not by itself particularly strong, in the absence of its human target. Integrating the results from both theory and experiments we dissect contributions to the mechanical stability of this protein.

## Introduction

Bacterial infections currently constitute a significant burden to human health, given that the diverse mechanism employed by these pathogens to infect the host are subject to continuous evolution. Furthermore, there is an increasing concern due to antibiotic resistance since it contributes to roughly 700,000 death annually^1^. Adherence is a crucial step in the bacterial infection process, allowing pathogens to colonize the host organism. Bacterial adhesins, a group of cell-surface proteins, utilize this adhesion capability to serve as essential virulence factors facilitating host colonization. These bacterial adhesins undergo physical stresses such as fluid flow, and understanding how bacteria respond to these mechanical cues is key^2,3^.

In the case of *S. aureus*, a Gram-positive bacterium well-known for causing a wide range of infections in both hospitals and community settings^4,5^, a special class of these adhesins are known as MSCRAMMs^6–10^. One such member of this family is ClfA a multidomain protein characterized by its immunoglobulin-like structure^11–13^. ClfA can be partitioned into two discernible regions: the R region and the A region (Fig. 1a). The R region, also known as Sdr, primarily comprises the dipeptide combination of aspartate and serine. The A region, in turn, exhibits further sub-division into three distinctive domains – N1, N2, and N3. It is noteworthy that the N1 domain lacks structural order, while the N2 and N3 domains are responsible for binding to a specific fragment of Fg. This binding is proposed to take place via the “dock lock and latch” (DLL) mechanism^14,15,7,16^. In the context of the DLL mechanism, initial engagement occurs as the C-terminus of the gamma chain of Fg is securely anchored (dock), subsequently undergoing confinement (lock) between the N2 and N3 domains, through a process known as beta complementation (latch)^16^. This mechanism is not exclusive to ClfA but is also observed in various homologous proteins including Fibronectin Binding Protein A (FnBPA)^21,22^ The DLL mechanism is key, for instance, in the development of endocarditis, a severe and potentially life-threatening condition characterized by inflammation of the endocardium^17^. Endocarditis starts when *S. aureus* binds to the heart valves through ClfA, amongst other MSCRAMMS. The strength of the attachment is crucial to secure infection^18,3,16,19,20^.

**Figure 1.**
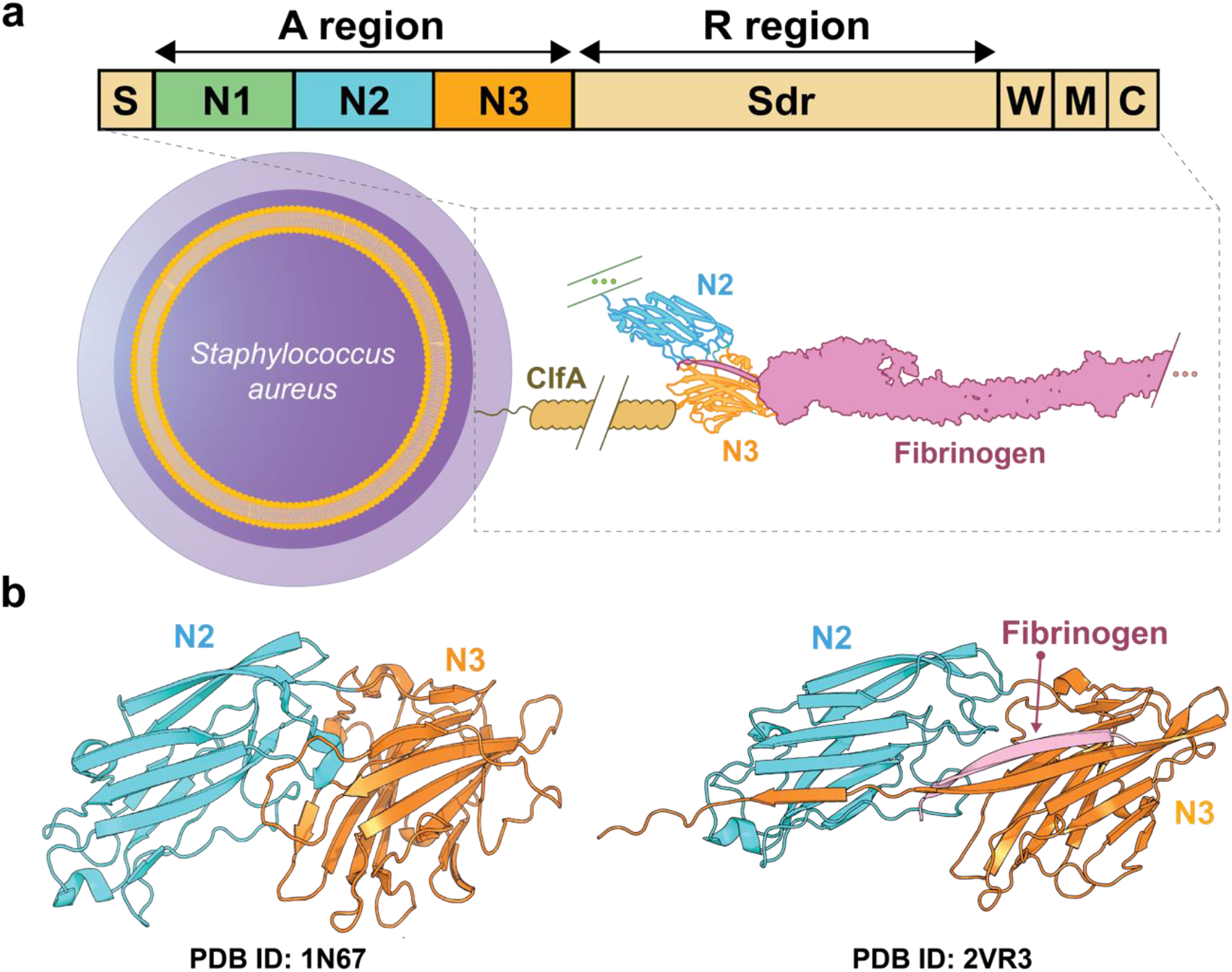
ClfA Structure and Attachment. (a) A schematic representation of ClfA attached to its target, with a diagram outlining the various domains that constitute ClfA. (b) ClfA structural changes upon fibrinogen binding: Comparison of the ClfA structure before and after binding to Fg through the DLL mechanism. PBD: 1N67 (-Fg) and PDB: 2VR3 (+Fg). N2 domain (orange), N3 domain (blue), and Fg (pink).

In a recent study, single molecule force spectroscopy (smFS) and atomistic molecular dynamics (MD) simulations have been used to study the mechanical robustness of the binding between Fg and a set of MSCRAMMs^16,23^. Although most of the work focused on SdrG, results were also presented for a few of its homologs, including ClfA. This study illustrated how MSCRAMMs bound to ligands can only be separated by strong forces exceeding 2 nN, rendering them as mechanically resilient as covalent bonds. This is remarkable considering the moderate (e.g. 5.8 μM) binding affinities of Fg peptides to MSCRAMMs^13^. It is not clear, however, whether the high mechanical strength is an evolved feature of the MSCRAMMs by themselves or only of their interactions with their ligands in the particular orientation probed in the smFS experiments on the protein constructs^24–27^.

In this article, we explore the intrinsic mechanical strength of ClfA. We use smFS to probe the mechanical unfolding of the protein and find that ClfA is not particularly strong in the absence of its human target. In particular, the N2 domain has a moderate mechanical strength, while the N3 domain is mechanically labile, resulting in a feature-rich and complex mechanical unfolding. We employ molecular simulations using a coarse-grained (CG) model to interpret the experimental results. Furthermore, combining a biophysical binding assay and smFS experiments in the presence of the soluble form of Fg, we find that the ligand by itself has no effect on ClfA’s mechanics.

## Methods

### Protein Expression and Purification

The gene encoding the chimeric polyprotein (191)_2_ - ClfAN2N3-(I91)_2_ was synthesized, and codon-optimized for efficient expression in *Escherichia coli* cells. For protein expression, the gene was cloned into pQE-80L vector (GenScript) and transformed onto *E. coli* BL21 (DE3) cells (Novagen). The transformed bacteria were grown in 750 mL of LB medium at 37 ºC until reaching an optical density at a wavelength of 600 nm (OD_600nm_). To induce protein expression, 1M of Isopropyl β-D-1-thiogalactopyranoside (IPTG) was added to the medium, and the culture was incubated overnight at room temperature with agitation at 180 rpm. After protein induction, the bacterial culture was collected by centrifugation at 4,000 x g and 4 ºC for 20 minutes. The resulting pellet was resuspended in 20 mL of Extraction Buffer (50 mM sodium phosphate monobasic, 50 mM sodium phosphate dibasic, and 300 mM NaCl – pH 7) supplemented with 160 µL of protease inhibitor cocktail (Thermo Scientific). To release the soluble protein into the media, mechanical disruption was performed using a French Press. The cell debris was then separated from the soluble fraction containing the proteins by centrifugation at 33, 000 x g for 30 minutes at 4 ºC. The supernatant was filtered using a filter of 0.22 µm (sartorius). The filtered supernatant was then incubated at 4 ºC with gentle shaking at 5 rpm for 1 hour in the presence of 5 mL of HisPur Cobalt Resin (GE Healthcare). After multiple rounds of resin cleaning using Extraction Buffer supplemented with 5 mM imidazole, the bound protein was eluted from the HisPur Cobalt Resin using the Elution Buffer (50 mM sodium-phosphate monobasic, 50 mM sodium-phosphate dibasic, 300 mM NaCl, 500 mM imidazole – pH 7). To further enhance the purity and remove any remaining contaminants, an additional purification step was conducted using size exclusion chromatography. Protein elution was carried out using a buffer containing 10 mM HEPES, 150 mM NaCl, 150 mM EDTA – pH 7.2. Following the size exclusion chromatography step, the purified protein sample was concentrated using Amicon 30 kDa ultrafiltration filters (Milipore). The concentrated protein sample was estimated to have a final concentration of 1 mg/mL. The purified samples were promptly snap-frozen in liquid nitrogen and stored at –80 ºC.

### Single-Molecule Force Spectroscopy

We have performed smFS experiments in the force-extension and force-ramp modes using a commercial Atomic Force Microscope (AFM) from Luigs and Neumann. 50 µl of a 5 µM polyprotein solution in 10 mM HEPES, 150 mM NaCl, 1 mM EDTA - pH 7.2 was deposited onto 15-mm-diameter coverslips coated with 10 nm gold and 25 nm titanium layer. Silicon nitride MLCT cantilevers from Bruker with a reflective 50 nm gold coating on their back side were used. Typical spring constants were in the range of 10-60 pN·nm^-1^ for force-extension and 5-20 pN·nm^-1^ for force-ramp. These cantilevers were calibrated using the thermal fluctuations method. Custom-written software in Igor Pro 6.37 (Wavemetrics) was employed for the collection and analysis of traces. All experiments were performed at room temperature.

### Molecular Dynamics Simulations

We have run molecular simulations of the force-induced unfolding of ClfA using the all-atom structure-based coarse-grained model by Whitford and co-workers^28,29^. Briefly, the model is coarse-grained to the level of protein heavy atoms, and its potential energy function includes native-centric harmonic terms for bonds and angles, a Fourier series for the torsions, a Lennard-Jones potential for non-bonded interactions between heavy atoms that are in contact in the native state and excluded volume interactions for all other pairs of atoms. Parameters were generated automatically from the coordinates of the crystal structure of ClfA (PDB ID: 1n67) using the SMOG webserver using default parameters. We calibrate the simulation temperature by comparing the fluctuations (RMSF) in the CG model from equilibrium runs at multiple temperatures with those from all-atom simulations in explicit water at 300 K for the same system. The atomistic runs were performed using the optimized CHARMM36m force field along with the tip3p model^30^. An octahedral box was utilized for the simulation setup. Upon solvation, the net charge of the system was neutralized. Subsequently, 100 ns energy minimization preceded a 1 µs production simulation, capturing system dynamics and interactions. At the chosen temperature (T*=0.7, in reduced units, SI Fig. S5) we run pulling simulations applying an external force to the protein ends at a range of pulling speeds. Ten simulation trajectories were run at each set of conditions. Simulation trajectories were analyzed using in-house Python scripts.

### Binding Assays by Microscale Thermophoresis

The (I91)_2_-ClfAN2N3-(I91)_2_ polyprotein was fluorescently labeled by utilizing the commercial protein Labeling Kit RED-NHS 2^nd^ Generation (Nanotemper Technologies GmbH, Munich, Germany). The dye carries a reactive NHS-ester group that reacts with primary amines in the protein to form a covalent bond. To label the protein, a concentration of 6.16 µM in a solution of 130 mM sodium bicarbonate, 50 mM NaCl – pH 8.2 was prepared. The protein was then incubated with a 3-fold excess of the fluorescent dye for 30 minutes at room temperature in absence of light. Any unreacted dye was eliminated by gel filtration column provided in the kit. The degree of labeling was 1.25 (dye/protein), based on the ratio between absorption at 280 nm and 650 nm. For the binding assay, unlabeled Fg was titrated into a fixed concentration of 34 nM of fluorescent (I91)_2_-ClfAN2N3-(I91)_2_. The experiment was conducted in PBS (150 mM NaCl, 50 mM phosphate pH 7.4) supplemented with 0.025% Tween® 20. Samples were loaded into capillaries and analyzed using a Monolith NT.115 system (Nanotemper Technologies GmbH, Munich, Germany) with an excitation power of 20% and 40% MST power. Data analysis was performed using the Nanotemper analysis software (MO. Affinity Analysis).

## Results

### Single-molecule Force Spectroscopy of ClfA using force-extension mode

We employed smFS by AFM to investigate the mechanical properties of N2 and N3 domains. We first produced the polyprotein (I91)_2_-ClfAN2N3-(I91)_2_, which consists of the N2 and N3 domains of ClfA flanked by dimeric handles comprised of the I91 domains from human cardiac titin. These I91 domains were included as fingerprints due to their well-documented mechanical properties in smFS studies^31–34^. To immobilize the protein onto the gold substrate, we introduced two cysteines at the C-terminus of our construct, facilitating its attachment through physical absorption. Single polyproteins were captured by gently pressing the cantilever onto the surface (Fig. 2a). Two different modes, namely force-extension and force-ramp were employed for the smFS experiments.

**Figure 2.**
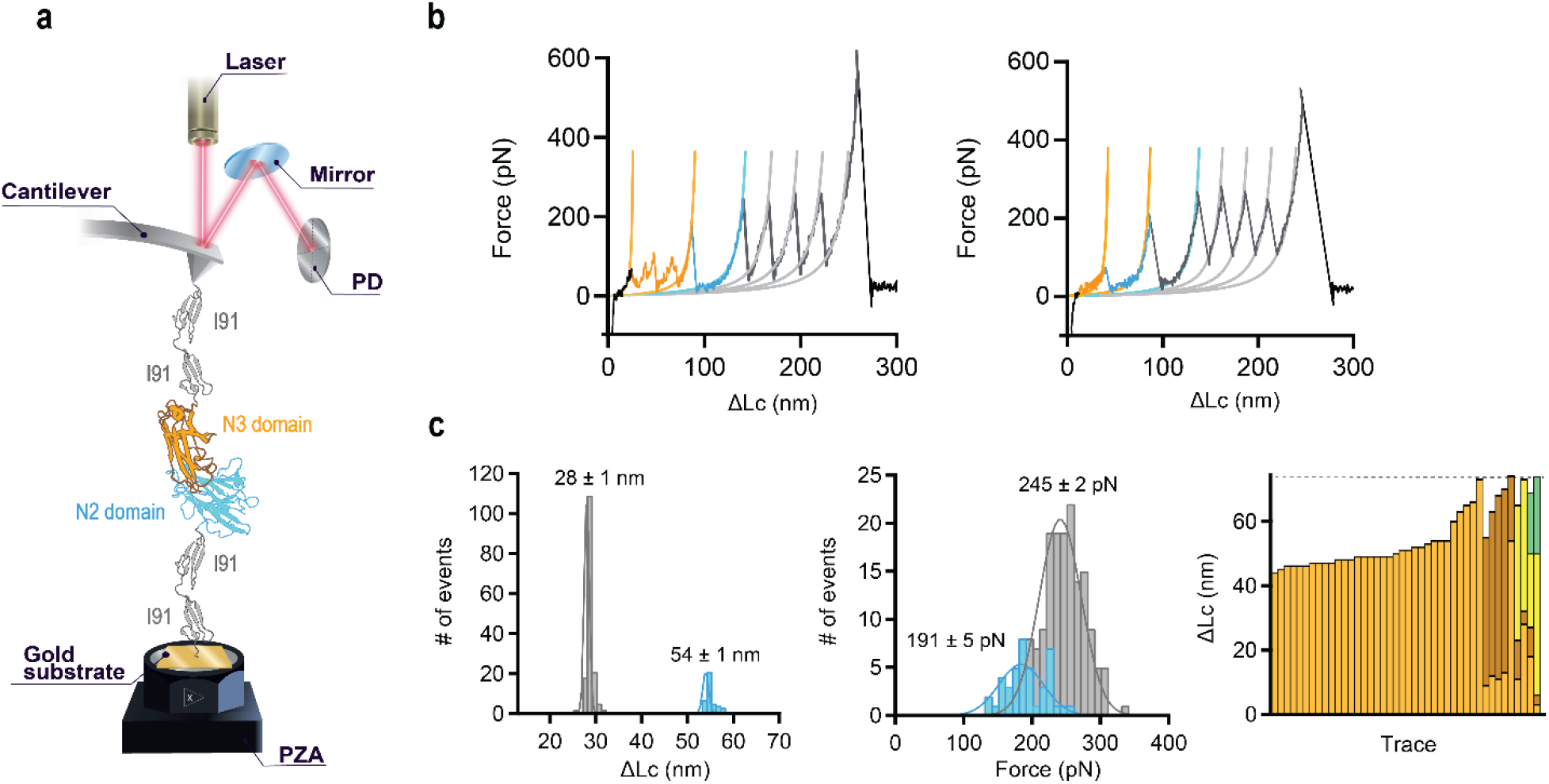
smFS analysis of (I91)_2_-ClfAN2N3-(I91)_2_ unfolding in force-extension. (a) Diagram illustrating the polyprotein (I91)_2_-ClfAN2N3-(I91)_2_ positioned between a cantilever tip and a gold surface. (b) Traces representing the two different unfolding patterns for the polyprotein sample. Lines represents fits to the Worm-like chain (WLC) model. c) Results obtained from force-extension mode experiments (N=42): Distribution plots depicting unfolding length and forces for I91 and N2 domains. The average step size (mean ± SEM) is 28 ± 1 nm for I91 and 54 ± 1 nm for N2. The average unfolding force (mean ± SEM) is 245 ± 2 pN for I91 and 191 ± 5 pN for N2. Additionally, unfolding length of N3 domain intermediates is presented, with different colors representing distinct numbers of intermediates. The dashed gray line indicates the theoretical extension of the N3 domain (73 nm). I91 domains (grey), N2 domain (blue), and N3 domain (orange).

In Figure 2b-c we present the results of the pulling experiments using the force-extension mode. In these experiments, the polyprotein was stretched at a constant speed of 400 nm·s^-1^ resulting in a trace with a characteristic sawtooth pattern, where each of the peaks corresponds to the unfolding of a single domain (Fig. 2b). The criterion for trace selection was the observation of rupture events of typical extension for at least three of the four titin domains, which appear late in the trace, just before the detachment peak at forces of 500-600 pN. In the resulting dataset (N=42) we observe a clear sequence of events, with ClfA unfolding first and the titin fingerprints appearing later in the traces (Fig. 2b). The assignment of the ClfA unfolding is however not unambiguous. We can clearly identify a peak in the distribution of extensions at contour a length of 54 ± 1 nm with unfolding force of 192 ± 4 pN, which we attribute to the N2 domain, with a theoretical extension of 55 nm (Fig. 2c). However, we do not find a clearly discernible pattern of extensions for the N3 domain. In fact, in 33 out of the 42 selected traces, we captured only a unique noisy unfolding event characterized by featureless initial rupture with a very low force, while in the remaining subset of traces, we found between two and four low force and short extension rupture events (representative examples are shown in Supplementary Fig. 1). When these events could be resolved, we obtained cumulative extensions that reach values comparable to the theoretical extension of 73 nm (Fig. 2c), but many times fall below that value due to the difficulty to measure the low force rupture events with our instrument. These findings suggest that the N3 domain is much more labile than N2.

### Single-molecule Force Spectroscopy in the ClfA using Force-ramp Mode

To investigate the unfolding of the N3 domain in further detail, experiments were conducted using a force ramp mode. In the force-ramp experiments, a progressively increasing force is applied to the polyprotein raising the force steadily with time^35–37^. This force ramp is achieved by implementing a feedback curve that dynamically corrects the position of the probe in response to the interaction occurring between the tip and the sample which allows to better control of the force applied. As the force gradually increases, the polyprotein undergoes mechanical deformations, resulting in the successive unfolding of its individual domains. The acquired force ramp data exhibit a distinctive staircase pattern, wherein each step corresponds to the unfolding event of a specific domain. In our smFS experiments, we used two distinct loading rates: an initial gentle ramp with a rate of 10 pN/s to facilitate the unfolding of both N2 and N3 domains, followed by a sharper ramp of 100 pN/s to induce the unfolding of the titin domains (Fig. 3).

**Figure 3.**
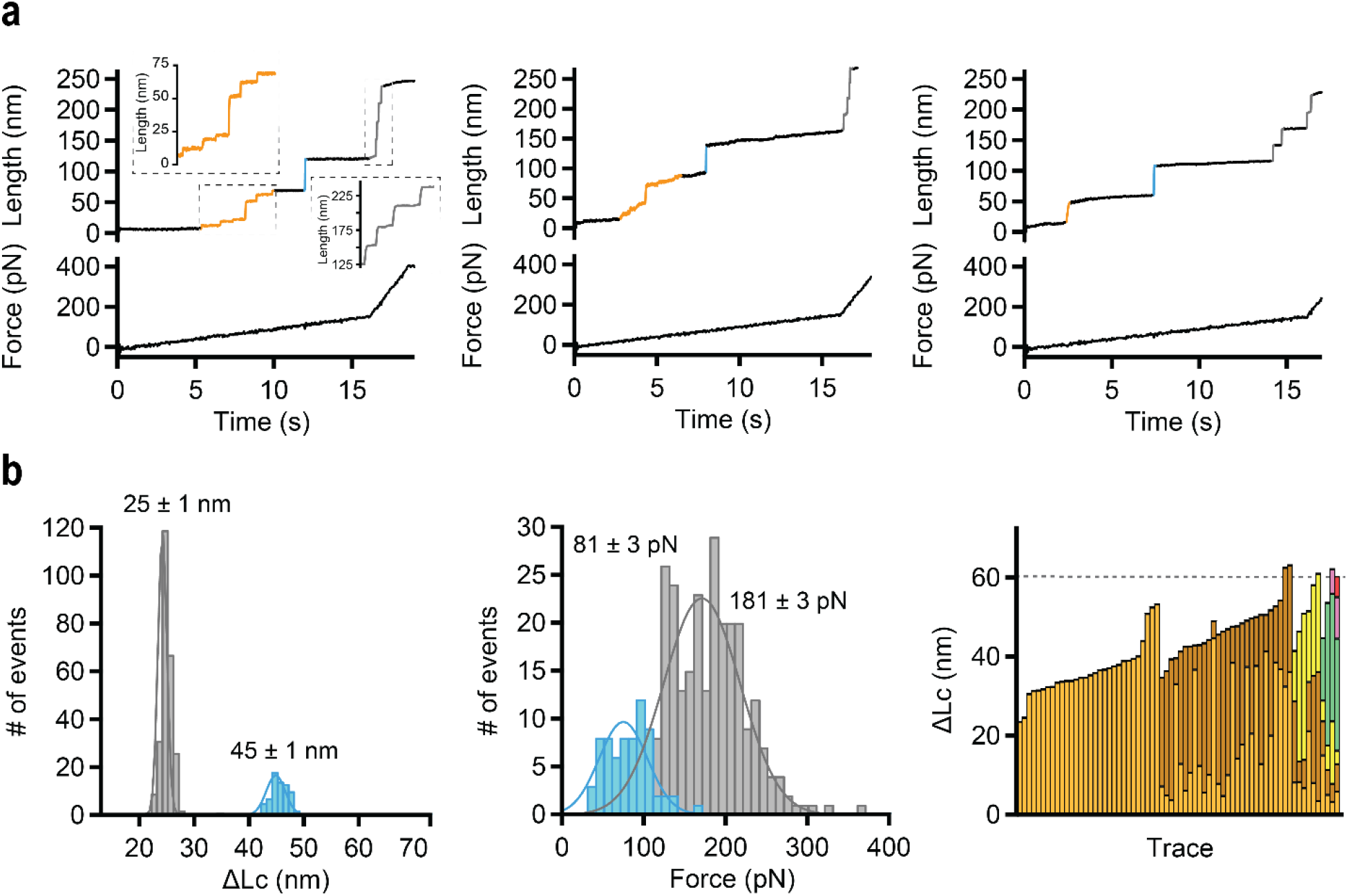
smFS analysis of (I91)_2_-ClfAN2N3-(I91)_2_ unfolding in force-ramp mode. (a) Traces representing the three different unfolding patterns for the polyprotein sample. b). Results obtained from force-ramp mode experiments (N=68): Distribution plots depicting unfolding length and forces for I91 and N2 domains. The average step size (mean ± SEM) is 25 ± 1 nm for I91 and 45 ± 1 nm for N2. The average unfolding force (mean ± SEM) is 181 ± 3 pN for I91 and 81 ± 3 pN for N2. Additionally, unfolding length of N3 domain intermediates is presented, with different colors representing distinct numbers of intermediates. The dashed gray line indicates the theoretical extension of the N3 domain (60 nm). I91 domains (grey), N2 domain (blue), and N3 domain (orange).

From these experiments, we kept a total of 68 traces that fulfilled our selection criteria. As found in the force-extension mode, the mechanical unfolding of the N2 domain results in a predictable jump in the extension of 45 ± 1 nm, associated to unfolding forces of 81 ± 3 pN. The traces are again more complex when we examine features associated with the unfolding of the N3 domain, which we can group in three distinct patterns (Fig. 3a). In seven traces, well-defined intermediates within the N3 domain are observable achieving the theoretical calculated extension of 60 nm. In contrast, most cases show intermediates with an unclear profile, resulting in a significantly varied unfolding pattern, characterized by extensions ranging from 40 to 55 nm. Finally, nine traces exhibit a single event of approximately 35 nm of extension. Interestingly, in one trace, a single event corresponding to the unfolding of both domains at a force of 48 pN was observed ( SI Fig. 2). Overall, we observe a mechanical unfolding complexity not common in proteins that play a mechanical role in their biological function. This confirms that the N3 domain, in the absence of Fg, is rather labile and without a well-defined unfolding pathway.

### Coarse grained simulations support a hierarchical unfolding pattern

Our findings so far reveal a hierarchy of unfolding events. N2 demonstrates mechanical properties consistent with previous findings on immunoglobulin-like domains, while N3 is highly labile, reaching the limits of resolution of our single-molecule instrument. To clarify these observations, we have run simulations using a structure-based, coarse grained simulation model that has been successfully used in the past to study force-induced unfolding of proteins^28,38–40^ (Methods). We first calibrated the model by comparing the fluctuations from equilibrium runs at multiple model temperatures with those observed using atomistic molecular dynamics at room temperature in explicit water (Methods and SI, Fig S5). At the chosen temperature, we have run simulations pulling from the protein termini at a constant extension rate (Fig. 4a). The force-extension curves reveal a pattern very similar to that found in the experiments. First, multiple low-force rupture events take place, affecting predominantly the N3 domain and interdomain interactions between N2 and N3. Only after the complete unfolding of N3, the N2 domain unfolds cooperatively, resulting in a single peak in all our simulation trajectories.

**Figure 4.**
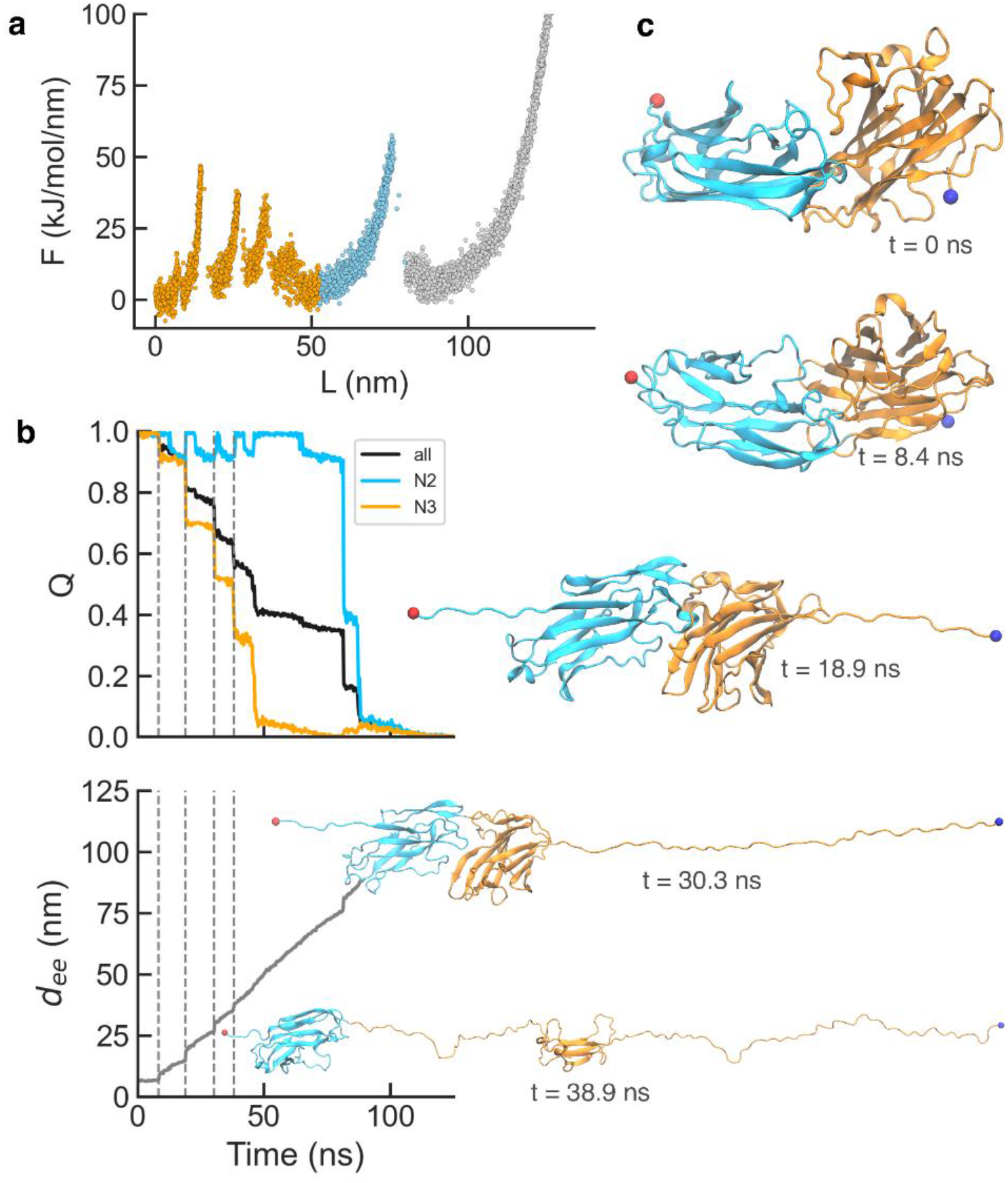
Coarse-grained molecular simulations of ClfA unfolding. (a) Overlay of the force-extension curves of ten independent trajectories. (b) Time series data for the fraction of native contacts, Q (top), and the end-to-end distance, d_ee_ (bottom), of a representative pulling trajectory. We show values for all the intramolecular contacts (black) and intradomain contacts within N2 (blue) and N3 (orange). Dashed vertical lines mark early local unfolding events primarily within N3. (c) Snapshots for selected time stamps marked with dashed lines in panel (b).

To gain further detail on the simulation results, we show the projection of a simulation trajectory on a relevant reaction coordinate for folding, the fraction of native contacts (*Q*) calculated both for the whole of the protein and each of the individual domains (Fig. 4b). We also show the end-to-end distance (*d*_ee_), which tracks the effect of the force in protein extension, and as expected from the simulation setup, changes linearly as a function of time. We find that unfolding of the N3 domain takes place in multiple steps, starting from the G’ strand (Fig. 4c), which in the Fg-bound state complements the E strand of N2. In the case of N2, the only early event is the rupture of a few contacts formed by a short N-terminal segment, and unfolding occurs later in the simulation trajectory and much more cooperatively (Fig. 4b). Despite some variation in the pattern of steps in N3 unfolding, all our simulation trajectories are consistent with a mechanism where different strands of N3 gradually peel off first from the immunoglobulin-like fold (Fig. 4c and SI Fig. S6). To test the robustness of our results, we have validated the proposed mechanism for two pulling speeds and found the same qualitative trends (SI Fig. S6 and S7), further asserting the validity of the proposed mechanism derived from the simulations. These observations recapitulate the experimental results.

### Binding of N2 and N3 Domains with Fibrinogen

In previous work by Bernardi and Gaub’s laboratories, ClfA was found to bind to the C-terminus of the gamma chain of Fg with a strength requiring a force of approximately 2 nN, establishing that the labile protein we study here forms, paradoxically, one of the most mechanically stable protein complexes described up to date. We have wondered whether the binding to Fg alters in some way the intrinsic mechanical strength of ClfA. To address this question, we have conducted smFS experiments in both force-extension (SI Fig. 4) and force-ramp (Fig. 5) modes, employing identical parameters as in the previous experiments, at 6.51 µM ClfA polyprotein concentration but now in presence of the Fg peptide in excess at a 3:1 peptide:protein ratio. In these conditions, Fg is expected to bind (I91)_2_-ClfAN2N3-(I91)_2_ construct given the *K*_D_ of 5.2 ± 0.5 µM, which we have determined from MST experiments (Methods and Fig. 5b). We note that this affinity is consistent with the dissociation constant previously published for ITC (*K*_*D*_ of 5.8 µM)^13^. In our smFS experiments in both modes, we observed similar unfolding patterns to those in the absence of Fg (Fig. 5a). Initially, the N3 domain unfolded in intermediates of varying lengths, followed by the unfolding of the N2 domain, and finally, the four titin domains. While all intermediates were detectable in some cases, they remained undetectable in many others. In the force-extension mode (SI Fig. 4), we measured an unfolding force for the N2 domain of 174 ± 4 pN, comparable to that obtained in the absence of Fg (Fig. 2a). This difference may be attributed to variability in the number of traces. In the force-ramp mode, the unfolding force of the N2 domain (81 ± 4 pN) matched that obtained in experiments without Fg (Fig. 5a). We hence conclude that ClfA is mechanically labile, regardless of the presence or absence of Fg, again due primarily to the very low resistance of the N3 domain.

**Figure 5.**
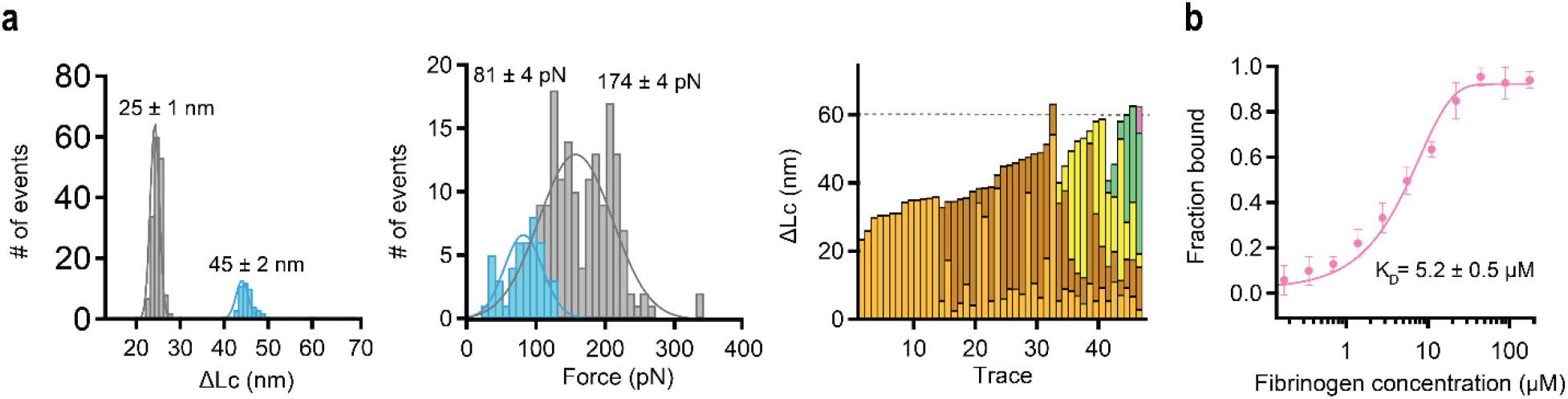
smFS analysis of (I91)_2_-ClfAN2N3-(I91)_2_ unfolding in presence of Fg. (a) Results obtained in force-ramp mode (N=46): Distribution plots depicting unfolding length and forces for I91 and N2 domains. The average step size (mean ± SEM) is 25 ± 1 nm for I91 and 45 ± 2 nm for N2. The average unfolding force (mean ± SEM) is 174 ± 4 pN for I91 and 81 ± 4 pN for N2. Additionally, unfolding length of N3 domain intermediates is presented, with different colors representing distinct number of intermediates. The dashed gray line indicates the theoretical extension of the N3 domain (60 nm). I91 domains (grey), N2 domain (blue), and N3 domain (orange). b) MST experiment to measure the binding between the Fg fragment and (I91)_2_-ClfAN2N3-(I91)_2_: K_D_ value of 5.2 ± 0.5 µM.

## Conclusion

The growing threat of bacterial infections and antibiotic resistance poses significant challenges to human health. MSCRAMMs, a class of cell surface proteins among *Staphylococcus* species, play a key role as virulence factors facilitating colonization of host organism by bacteria. These adhesins undergo physical stresses, and understanding how bacteria respond to these mechanical signals is important in addressing infections. Previous work has shown that these proteins bind extremely strongly to their partners in human cells, specifically to the Fg peptide, but it was not yet clear whether the mechanical strength was an evolved feature intrinsic to the protein or instead only determined by inter-protein interactions between Fg and the MSCRAMM.

Here, we have conclusively addressed this question. We have designed this study specifically to investigate the intrinsic mechanics of ClfA utilizing smFS, aided by molecular simulations using a coarse-grained model and MST. We have found that, regardless of the presence or absence of its binding partner, ClfA is mechanically labile. Our findings have unveiled a remarkable mechanical instability within ClfA’s binding region. Notably, while the N2 domain demonstrates sufficient strength, allowing it to unfold in a single event, the N3 domain exhibits substantial instability, presenting challenges in its precise characterization using smFS. Additionally, our research demonstrates that the robust binding between ClfA and its ligand results from their specific orientation. In this sense, it is well-known that the mechanical stability of proteins depends on the orientation of the applied force. In the present study we apply force to the N2N3 tandem following a force vector that connects the N- and C-termini. This orientation differs from the force vector that encounters the tandem *in vivo* when bound to Fg. In this case, the force propagates along the vector that connects Fg with N3 through the DLL mechanism (Fig 1b). What is surprising is the large difference in unfolding force in the presence and absence of ligand, which suggests that ClfA does not feature a high mechanically stable *per se*, and only the binding of Fg transforms it into one of the highest mechanical interactions ever reported in a protein. Indeed, the lability of the N3 domain seems to play a crucial role in facilitating the DLL mechanism, enabling the conformational shift of one of the G-strand toward the N2 domain, effectively enclosing Fg during the union. Overall, we reveal here new aspects on the mechanics of ClfA, a bacterial infection-related protein whose mechanical strength covers a force spectrum that goes from tens of pN to nN.

## Data availability

Data supporting the findings of this study are available from the corresponding author upon reasonable request.

## Supporting information

This article contains supporting information.

## Acknowledgements

This work has been supported by grants PID2019-109087RB-I00 to R.P.-J. from Spanish Ministry of Science and Innovation and Spanish Research Agency (MCIN/AEI). This work has received funding from the European Union’s Horizon 2020 research and innovation program under grant agreement No 964764 to R.P.-J. Financial support to D. D comes from Eusko Jaurlaritza (Basque Government) IT1584-22 and from the Spanish Ministry of Science and Universities through the Office of Science Research (MINECO/FEDER) through grant PID2021-127907NB-I00 from MCIN/AEI. A.L.C. acknowledges financial support from grants: PID2022-137977OB-I00 funded by MCIN/AEI/ 10.13039/501100011033. We acknowledge the Severo Ocho Excellence Program grant CEX2021-001136-S funded by MCIN/AEI 10.13039/501100011033. This work was performed under the Maria de Maeztu Units of Excellence Program from Q5, grant no. MDM-2017-0720 funded by MCIN/AEI.

