## Supplementary Material for "The Strongest Protein Binder is Surprisingly Labile"

### Sequences

#### $(I91)_2$-ClfAN2N3-$(I91)_2$ – Uniprot (Q53653)

MRGSHHHHHHLIEVEKPLYGVEVFVGETAHFEIELSEPDVHGQWKLKGQPLTASPDCEIIEDGKKHIL
ILHNCQLGMTGEVSFQAANAKSAANLKVKELLIEVEKPLYGVEVFVGETAHFEIELSEPDVHGQWKLK
GQPLTASPDCEIIEDGKKHILILHNCQLGMTGEVSFQAANAKSAANLKVKELRSRSVAADAPAAGTDI
TNQLTNVTVGIDSGTTVYPHQAGYVKLNYGFSVPNSAVKGDTFKITVPKELNLNGVTSTAKVPPIMAG
DQVLANGVIDSDGNVIYTFTDYVNTKDDVKATLTMPAYIDPENVKKTGNVTLATGIGSTTANKTVLVD
YEKYGKFYNLSIKGTIDQIDKTNNTYRQTIYVNPSGDNVIAPVLTGNLKPNTDSNALIDQQNTSIKVY
KVDNAADLSESYFVNPENFEDVTNSVNITFPNPNQYKVEFNTPDDQITTPYIVVVNGHIDPNSKGDLA
LRSTLYGYNSNIIWRSMSWDNEVAFNNGSGSGDGIDKPVVPEQPDEPGEIEPIPERSLIEVEKPLYGV
EVFVGETAHFEIELSEPDVHGQWKLKGQPLTASPDCEIIEDGKKHILILHNCQLGMTGEVSFQAANAK
SAANLKVKELLIEVEKPLYGVEVFVGETAHFEIELSEPDVHGQWKLKGQPLTASPDCEIIEDGKKHIL
ILHNCQLGMTGEVSFQAANAKSAANLKVKELSSCC

#### C-terminus of Fibrinogen Gamma Chain – Uniprot (P02679)

GEGQQHHLGGAKQAGDV

### Theoretical Extension for N2 and N3 Domains (PDB ID: 1N67)

| Force-extension |
| --- |
| **N2 domain** ⇢ 149 residues x 0.4 nm per residue – 4.63 nm folded = 54.97 nm |
| **N3 domain** ⇢ 190 resides x 0.4 nm per residue – 2.7 nm folded = 73.3 nm |
| **N2 and N3 domains** ⇢ 128.27 nm |

| Force-ramp |
| --- |
| **N2 domain** ⇢ 149 residues x 0.33 nm per residue – 4.63 nm folded = 44.54 nm |
| **N3 domain** ⇢ 190 resides x 0.33 nm per residue – 2.7 nm folded = 60 nm |
| **N2 and N3 domains** ⇢ 104.54 nm |

### Theoretical Extension for N2 and N3 Domains (PDB ID: 2VR3)

| Force-extension |
| --- |
| **N2 domain** ⇢ 149 residues x 0.4 nm per residue – 4.61 nm folded = 54.99 nm |
| **N3 domain** ⇢ 190 resides x 0.4 nm per residue – 6.39 nm folded = 69.61 nm |
| **N2 and N3 domains** ⇢ 124.6 nm |

| Force-ramp |
| --- |
| **N2 domain** ⇢ 149 residues x 0.33 nm per residue – 4.61 nm folded = 44.56 nm |
| **N3 domain** ⇢ 190 resides x 0.33 nm per residue – 6.32 nm folded = 56.38 nm |
| **N2 and N3 domains** ⇢ 100.94 nm |

### Supplementary Figures

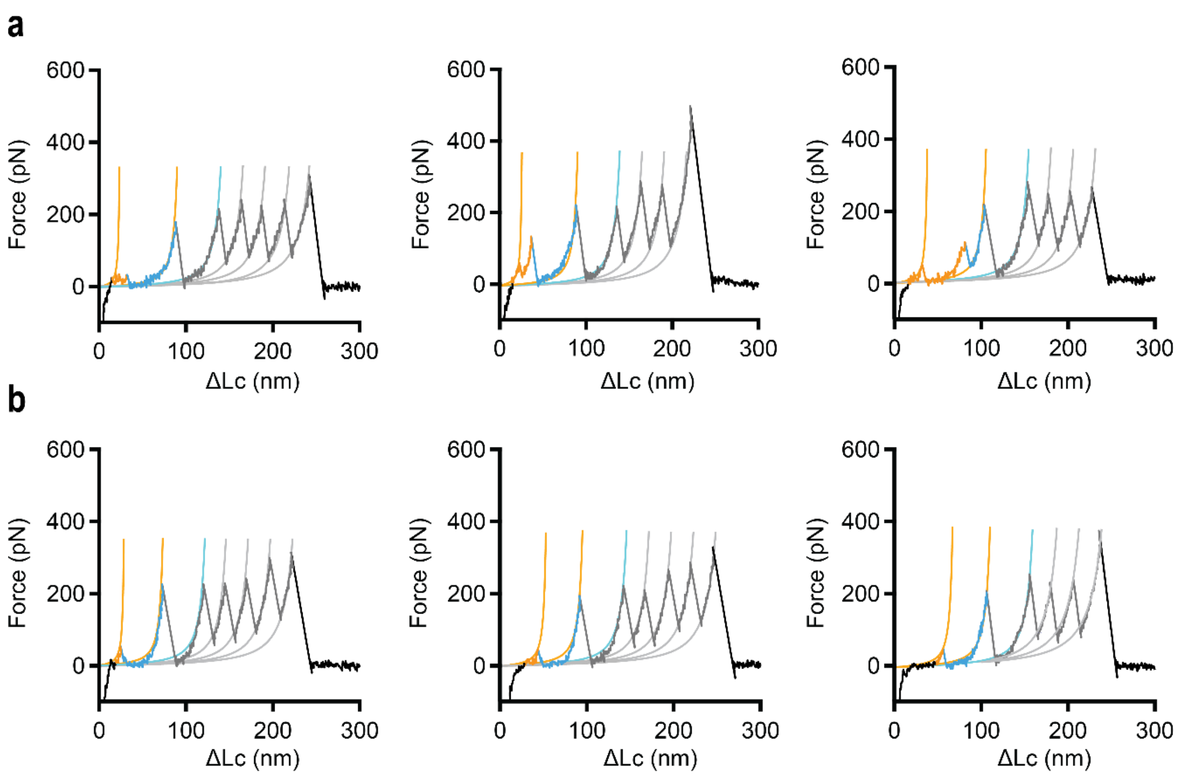


**Supplementary Figure S1. Representative smFS traces of (I91)$_2$-ClfAN2N3-(I91)$_2$ unfolding in force-extension mode**. a) Illustrative traces showcasing various rupture events within the N3 domain, with the summation of all N3 extensions aligning with the theoretical extension (73 nm). b) Exemplary traces illustrating the detection of a singular event within the N3 domain. I91 domains (grey), N2 domain (blue), N3 domain (orange). Traces represented following the Worm-like-chain (WLC) model.

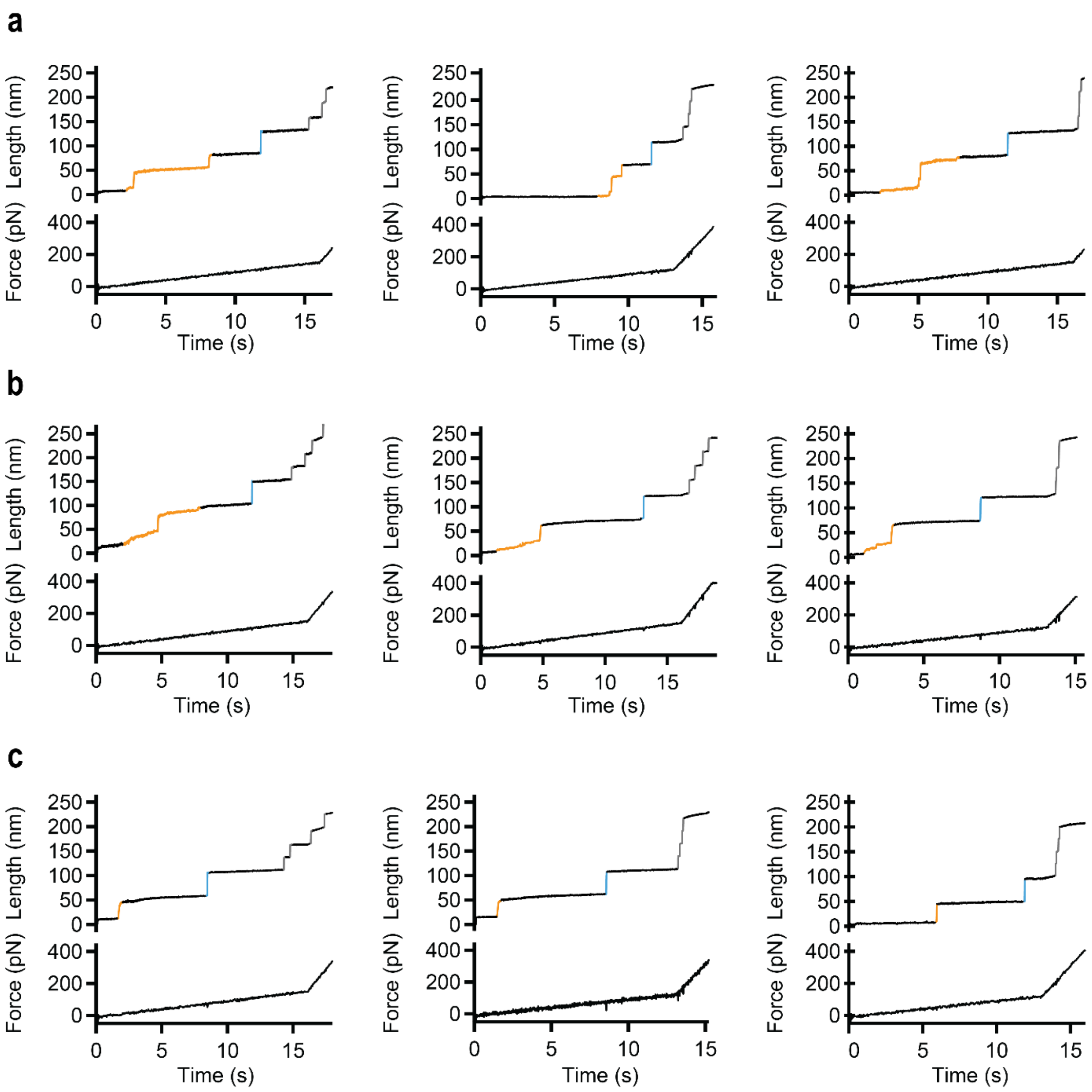


**Supplementary Figure S2. Representative smFS traces of (I91)$_2$-ClfAN2N3-(I91)$_2$ unfolding in force-ramp mode.** a) Illustrative traces showcasing various rupture events within the N3 domain, with the summation of all N3-extension aligning with the theoretical extension (60 nm). b) Illustrative traces where not all the N3 domain rupture events are well-defined. C) Exemplary traces illustrating the detection of a singular event within the N3 domain. I91 domains (grey), N2 domain (blue), N3 domain (orange).

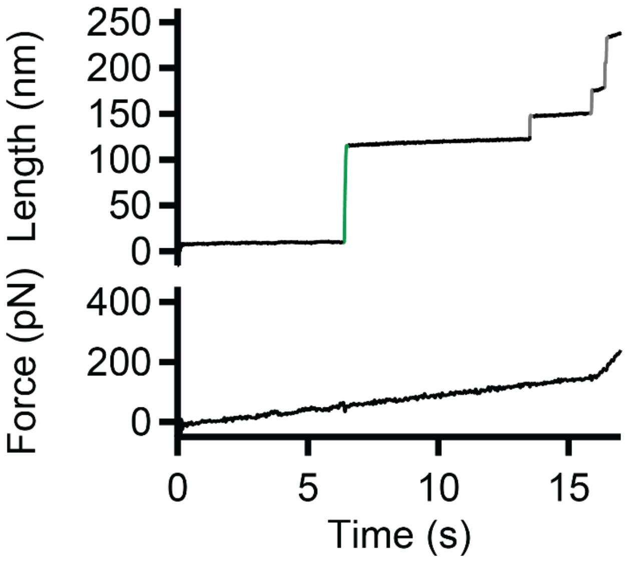


**Supplementary Figure S3. Representative smFS trace of $(I91)_2$-ClfAN2N3-$(I91)_2$ unfolding in force-ramp mode.** Trace illustrating the unfolding of N2 and N3 domains simultaneously, with the green color indicating the unfolding of both N2 and N3 domains.

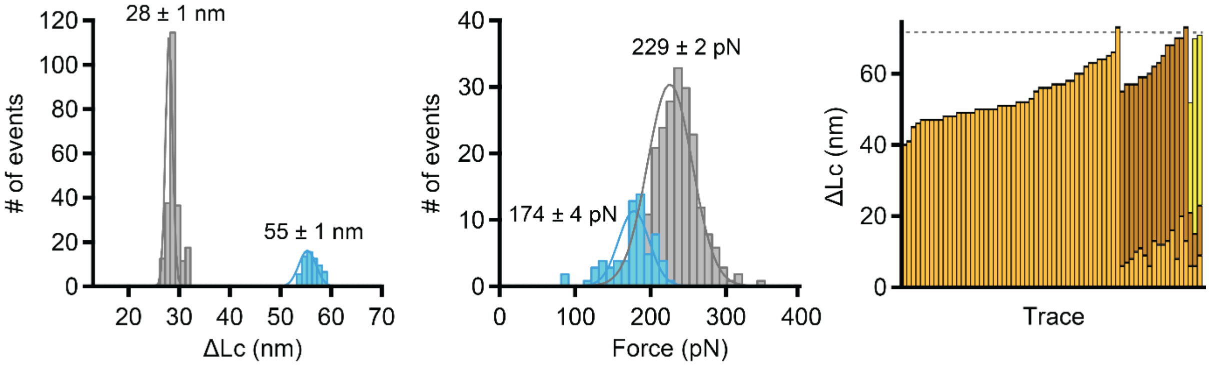


**Supplementary Figure S4. Representative smFS analysis of $(I91)_2$-ClfAN2N3-$(I91)_2$ unfolding in presence of Fg.** Results obtained from force-extension mode experiments (N=66): Distribution plots depicting unfolding length and forces for I91 and N2 domains. The average step size (mean ± SEM) is 28 ± 1 nm for I91 and 55 ± 1 nm for N2. The average unfolding force (mean ± SEM) is 229 ± 2 pN for I91 and 174 ± 4 pN for N2. Additionally, unfolding length of N3 domain intermediates is presented, with different colors representing distinct numbers of intermediates. The dashed gray line indicates the theoretical extension of the N3 domain (70 nm). I91 domains (grey), N2 domain (blue), and N3 domain (orange).

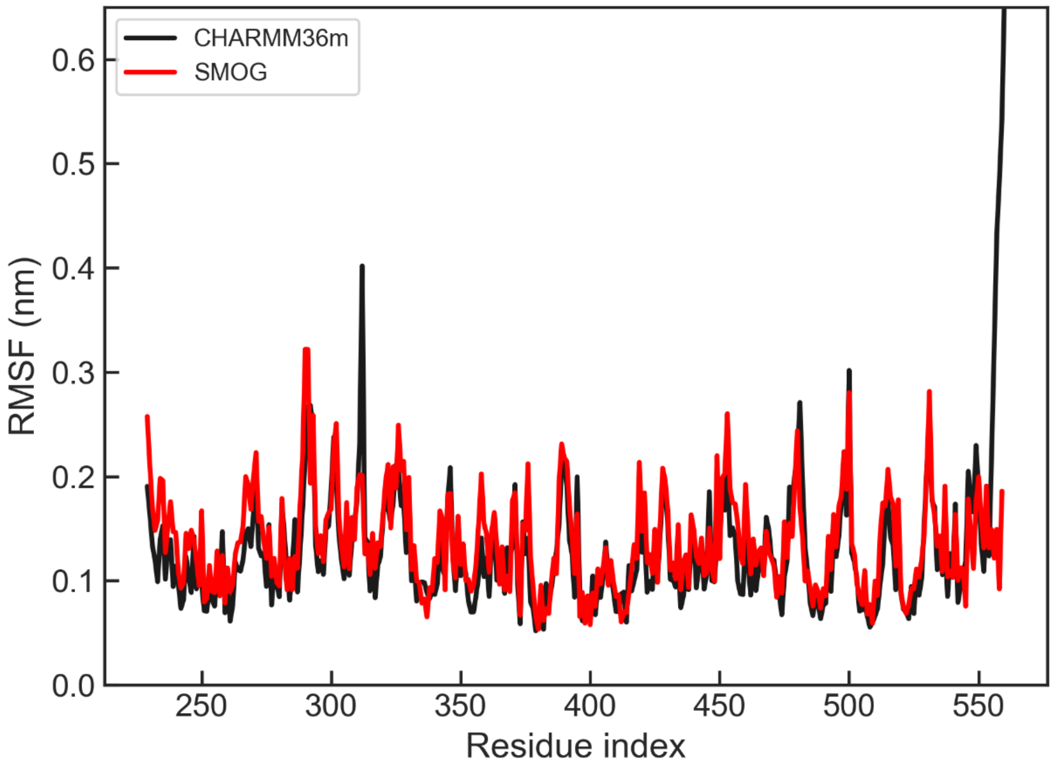


**Supplementary Figure S5. Calibration of the coarse-grained model.** Comparison between the RMSF calculated from an atomistic MD simulation in explicit solvent using the CHARMM36m force field (black) and that from the coarse-grained model at a reduced temperature of T*=0.7 (red).

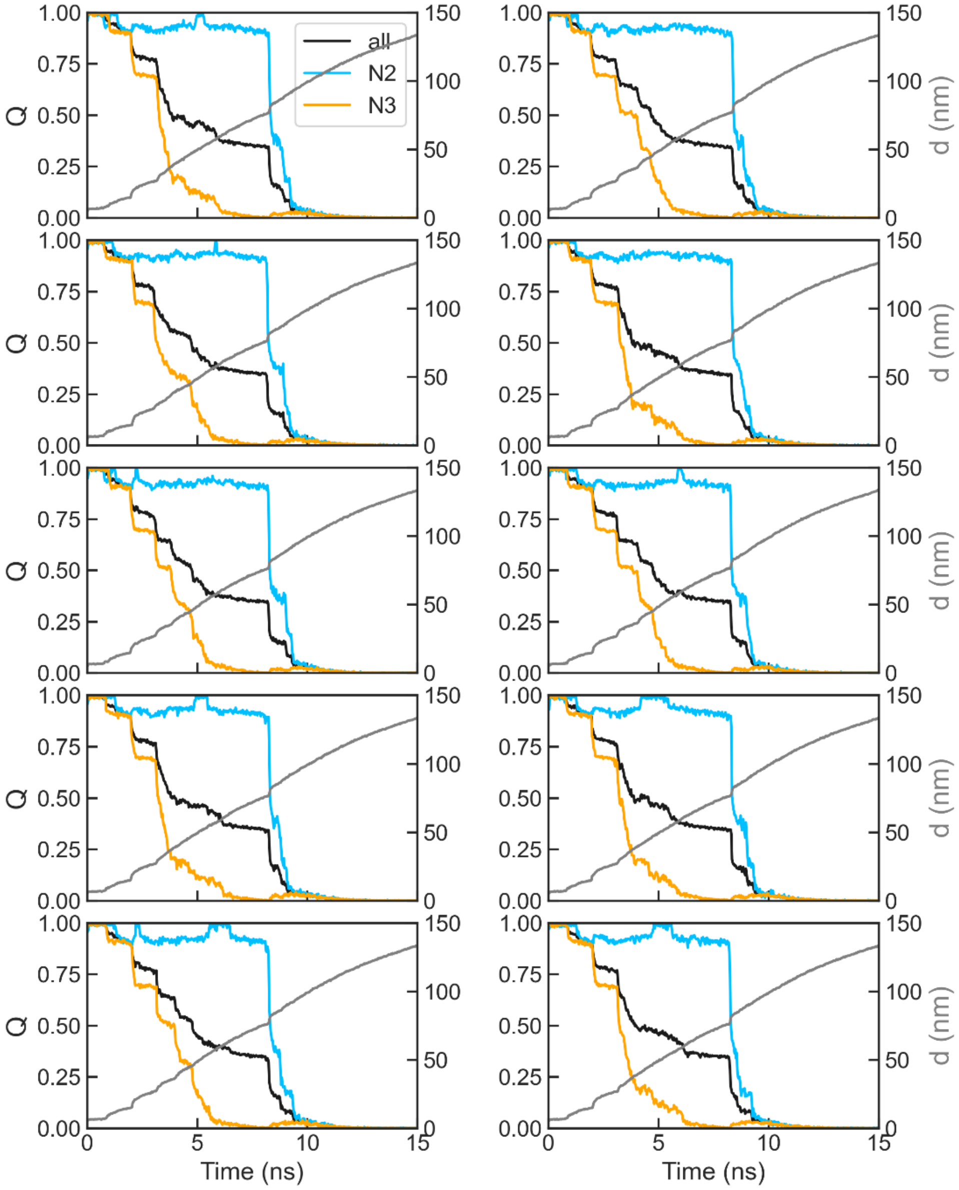


**Supplementary Figure S6. Individual simulation trajectories of mechanical unfolding at a pulling speed of 1e-2 nm/ps.** For each trajectory, we show time series data for the fraction of native contacts, Q (top), and the end-to-end distance, dee (bottom), of a representative pulling trajectory. We show values for all the intramolecular contacts (black) and intradomain contacts within N2 (blue) and N3 (orange).

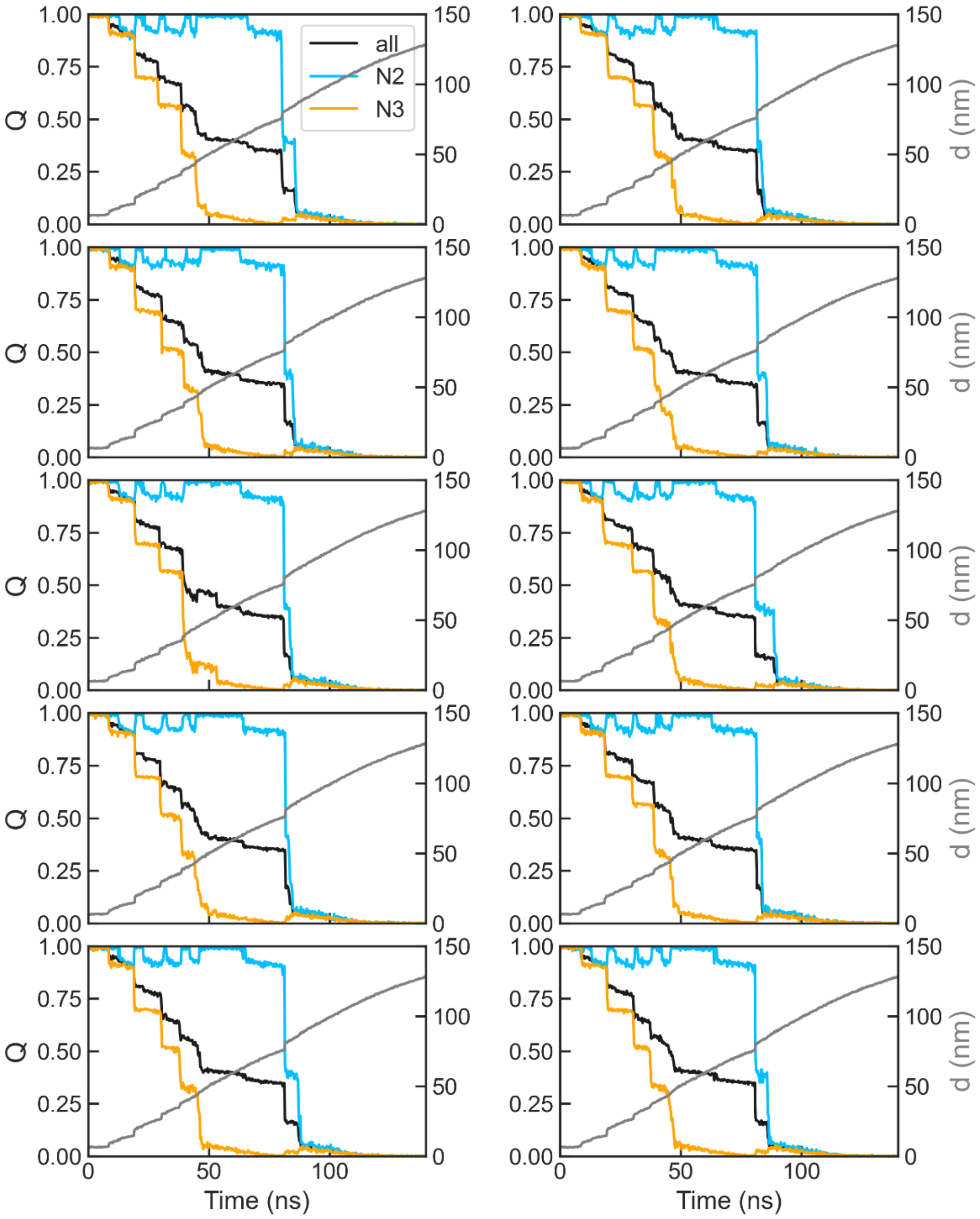


**Supplementary Figure S7. Individual simulation trajectories of mechanical unfolding at a pulling speed of 1e-3 nm/ps.** For each trajectory, we show time series data for the fraction of native contacts, Q (top), and the end-to-end distance, dee (bottom), of a representative pulling trajectory. We show values for all the intramolecular contacts (black) and intradomain contacts within N2 (blue) and N3 (orange).
